# Loss of RNF2 delays tumour development in BAP1-deficient mesothelioma

**DOI:** 10.1101/2024.04.26.591241

**Authors:** Nick Landman, Danielle Hulsman, Jitendra Badhai, Maarten van Lohuizen

## Abstract

The tumour suppressor gene *BAP1* is mutated in more than half of the malignant mesothelioma patients. This catalytic subunit of Polycomb repressive deubiquitinating (PR-DUB) complex, plays an important role in maintaining gene expression levels by deubiquitinating the PRC1-mediated histone H2A lysine 119 mono-ubiquitination (H2AK119ub1). Published studies report varying degrees of importance of H2AK119ub1 in Polycomb-regulated gene expression in different cell types. Recently published data by our own lab suggests a global redistribution of the H2AK119ub1 mark from promoter to intergenic regions upon loss of BAP1. PRC1-mediated mono-ubiquitination is dependent on the E3 ubiquitin ligase function of RNF2 (RING1B). Here, by knocking-out *Rnf2*, we show that loss of H2AK119ub1 levels leads to a decrease in clonogenic potential of *Bap1*-deficient mesothelioma cells *in vitro* and a delay in tumour onset *in vivo*.

## Introduction

BAP1 is a ubiquitin C-terminal hydroxylase and is the catalytic subunit of the Polycomb repressive de-ubiquitinase complex (PR-DUB). This complex, via BAP1, deubiquitinates PRC1-mediated histone H2A lysine 119 mono-ubiquitination (H2AK119ub1) (1, 2, 3). BAP1 is a tumour suppressor that has been found to be mutated or deleted in multiple malignancies in a high frequency, in more than half of mesothelioma cases, uveal melanoma (43%), cholangiocarcinoma (25%), renal cell clear cell carcinoma (23%), and others (4, 5, 6, 7, 8). BAP1’s tumour-suppressive mechanism remains elusive however its DUB activity suggests an interplay with PRC1-mediated mono-ubiquitination of H2AK119. In embryonic stem (ES) cells it has been shown that the loss of BAP1 leads to an overall increase in H2AK119ub1 levels, titrating PRC2 away from its target genes (9). In a recently published study by our lab, we show that in mesothelioma cells loss of BAP1 results in an overall increase in levels of the PRC2-mediated H3K27me3 mark and in a shift of H2AK119ub1 from promoter to intergenic regions (10). Similar observations are made in uveal melanoma upon BAP1 loss (11). However, in mesothelioma transcription start site regions co-occupied by H2AK119ub1 and H3K27me3 gene expression seems to follow the PRC2-mediated mark. Additionally, a recent study in HAP1 cells suggests that transcriptional changes upon gain of H2AK119ub1 are partly independent of PRC2 (12).

PRC1-mediated mono-ubiquitination is dependent on the E3 ubiquitin ligase function of RNF2 (RING1B) and RNF1 (RING1A) which on its turn is required for the recruitment of PRC2 on chromatin leading to H3K27me3 deposition (2, 13). In mouse embryonic stem cells, fibroblasts, liver, and pancreatic tissue it is shown that BAP1 inactivation causes RNF2-dependent apoptosis. However, in melanocytes and mesothelial cells, BAP1 loss had little impact on survival. Further, this study suggested that loss of BAP1 only promotes tumorigenesis in cells with RNF2-independent apoptosis (14). To investigate the role of RNF2 in tumorigenic potential of *Bap1*-deficient mesothelioma in this study we generated *Rnf2* knock-outs in mesothelioma cell lines. We show that loss of the E3 ubiquitin ligase subunit RNF2 limits the clonogenic potential of *Bap1*-deficient, but not *Bap1*-proficient, mesothelioma cells and show that this leads to a delay in tumour development in xenograft models.

## Results

### Generating *Rnf2* knock-out mesothelioma cell lines

BAP1 is a deubiquitinase protein functioning in the PR-DUB complex removing the PRC1-mediated H2AK119ub1 mark. PRC1 catalyses monoubiquitination of H2A via the E3 ubiquitin ligases RNF1 (RING1A) and RNF2 (RING1B). Previously, we and others have shown that loss of BAP1 leads to an increase in H2AK119ub1 levels.

However, the precise consequence and role of PRC1-mediated ubiquitination on mesothelioma tumours formation remains inconclusive. Therefore, we have investigated the effect on tumorigenic potential of mesothelioma cells upon knock-out of *Rnf2*. In a recent study, we have derived cell lines from tumours isolated from autochthonous mouse mesothelioma models, referred to as NC (*Nf2*-/-, *Cdkn2ab*-/-) and BNC (*Bap1*-/-, *Nf2*-/-, *Cdkn2ab*-/-). These cell lines were transfected to stably express the Cas9 protein. We transiently transfected early passage cell lines (within the first 4 – 8 passages) with a pLKO plasmid with a guide RNA targeting *Rnf2* and a GFP construct, cells were then used to generate *Rnf2* knock-out clones (**Figure 1a**). We sorted the transfected cell lines by flow cytometry for their GFP signal and plated them out as single cells (**Figure 1b**). Surviving-single cell colonies were expanded and sent for sequencing to validate CRISPR/Cas9-mediated knock-out of *Rnf2*. For both the BNC and NC cell lines we were able to identify monoclonal colonies where the CRISPR/Cas9 construct has efficiently cut the DNA (**Figure 1c**). To validate whether these cell lines were true *Rnf2* knock-out we performed western blot analysis showing full knock-out for RNF2 protein in the BNC and NC lines.

**Figure 1.**
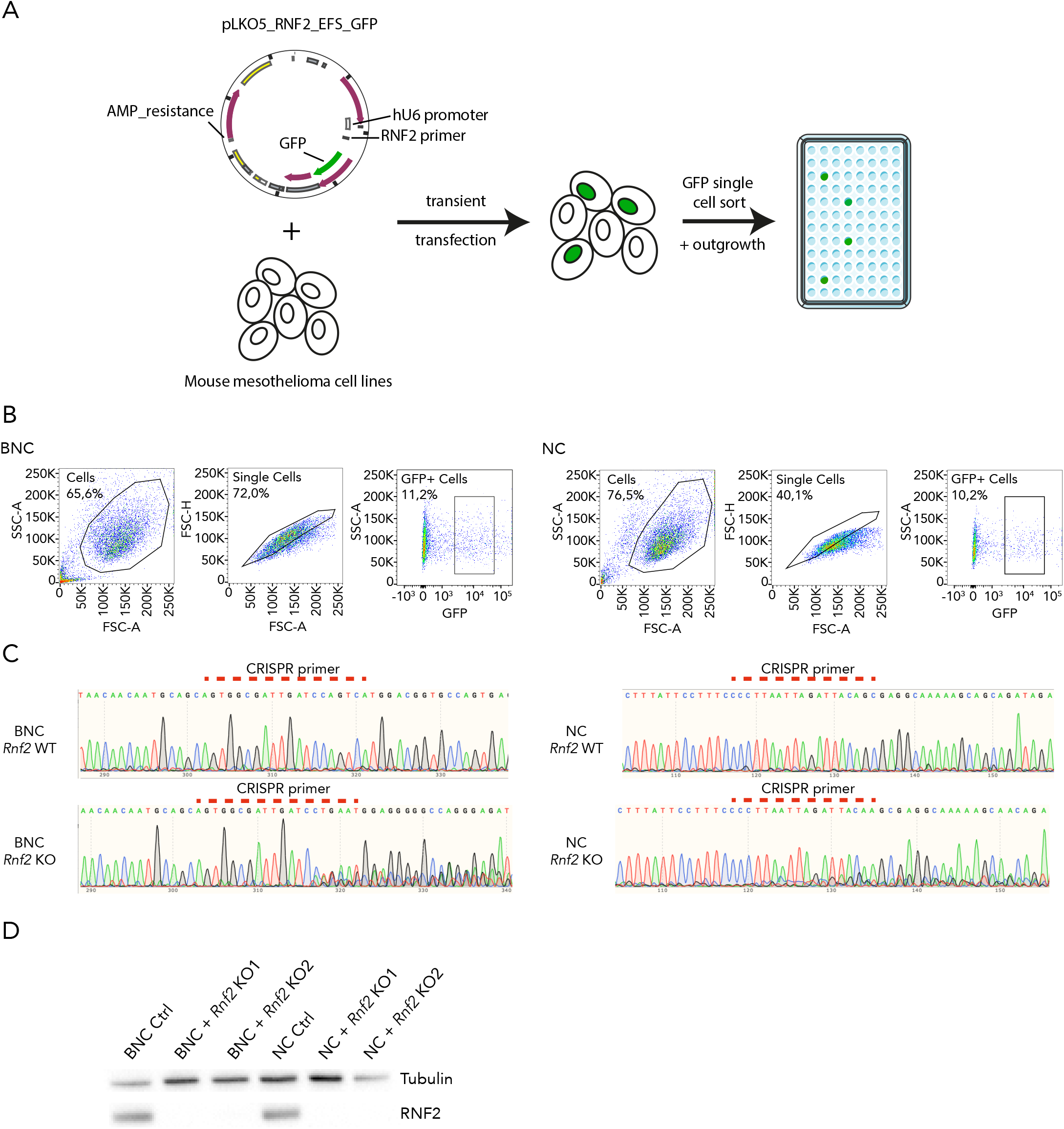
Generating *Rnf2* knock-out mouse mesothelioma cell lines. **A**, Schematic overview of experimental setup to obtain single cell *Rnf2* knock-out cells. **B**, Gating strategy of flowcytometry for single cell sorting of GFP positive cells. **C**, Targeting sequence of the used CRISPR guide RNAs and the altered sequence after Cas9 endonuclease activity in the *Bap1*-deficient (BNC) and proficient (NC) mesothelioma cell line. **D**, Western blot showing the total loss of RNF2 protein in the sorted single cell clones. Tubulin was used as a loading control.

### *Rnf2* knock-out does not influence cell morphology and proliferation but lowers the clonogenic potential of BAP1-deficient mesothelioma cells

As PRC1-mediated H2A monoubiquitination is widely implicated in cell differentiation and plasticity we checked whether knock-out of *Rnf2* led to massive changes in mesothelioma tumours cell phenotype. Neither in the *Bap1*-deficient nor the *Bap1*-proficient we were able to identify any changes in phenotype between *Rnf2* knock-out and *Rnf2* wild-type cells (**Figure 2a**). Analysing the proliferation rate of the generated *Bap1*-deficient *Rnf2* knock-out cell line did not show deviant proliferation compared to *Rnf2* wild-type cells. Similarly, *Rnf2* knock-out and wild-type *Bap1*-proficient cells also proliferate at a comparable rate (**Figure 2b**).

**Figure 2.**
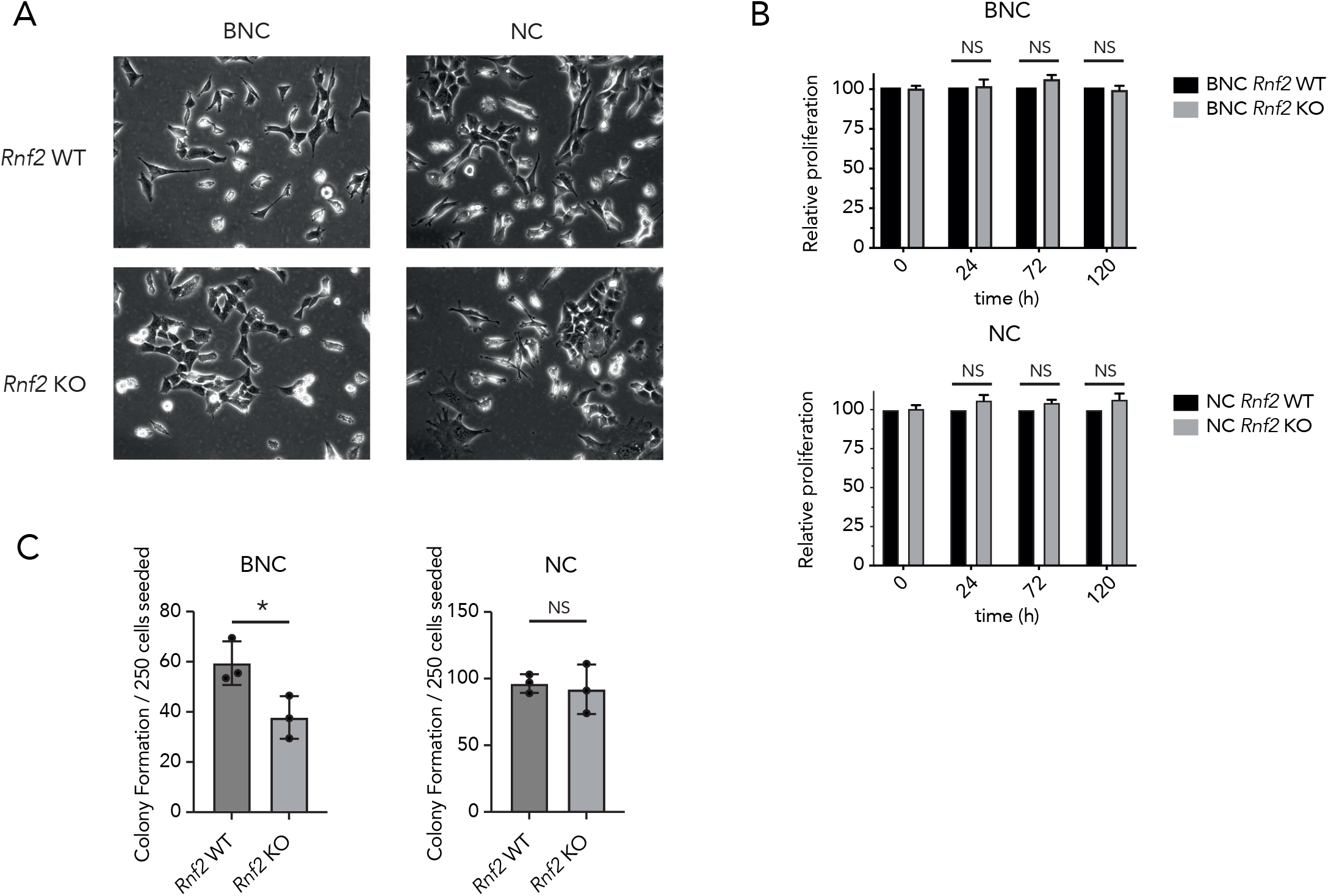
*Rnf2* knock-out does not influence cell morphology and proliferation but lowers the clonogenic potential of *Bap1*-deficient mesothelioma cells. **A**, Representative images of *Bap1*-proficient and deficient cell lines with and without *Rnf2* knock-out, these show no morphological differences upon loss of *Rnf2*. **B**, Relative proliferation rates of the cell lines show no difference between *Rnf2* knock-out and wild-type cell lines irrespective of the *Bap1*-status (mean ± s.d.; n = 3 independent experiments). **C**, Colony formation assays show that knock-out of *Rnf2* decreases the clonogenic potential of *Bap1*-deficient cells but not of *Bap1*-proficient cells (mean ± s.d.; n = 3 independent experiments). P values were determined by two-tailed unpaired Student’s t-test; *P < 0.05

Next, we analysed whether knocking-out *Rnf2* had an effect on the clonogenic potential of mesothelioma cells. In the *Bap1*-proficient NC line, we do not observe any significant difference in the amounts of colonies formed from a single cell. However, cells that were deficient for *Bap1* and knocked-out for *Rnf2* showed a significantly lower clonogenic potential when compared to *Rnf2* wild-type cells (**Figure 2c**).

### *Bap1*-deficient mesothelioma cells knocked-out for *Rnf2* have delayed tumour onset

To see whether the lower clonogenic potential of *Rnf2* knock-out *Bap1*-deficient mesothelioma cells is also reflected *in vivo*, we performed mouse xenograft experiments. Before injection of the cell lines into mice we analysed the expression of the PRC1 and PRC2-mediated chromatin marks, H2AK119ub1 and H3K27me3 respectively, in both the *Bap1*-proficient and deficient lines. In line with observations in previous studies, we observe lower overall H2AK119 monoubiquitination in the cells proficient for *Bap1*, indicative of BAP1’s deubiquitinase function (**Figure 3a**). Additionally, in both *Bap1*-proficient and deficient cells we observe lower H2AK119ub1 levels in *Rnf2* knock-out cells compared to *Rnf2* wild-type cells, which is in line with the ubiquitination function of the PRC1 complex. Surprisingly, the PRC2-mediated H3K27me3 mark levels are lower in *Rnf2* knocked-out *Bap1*-proficient cells but not in *Rnf2* knocked-out *Bap1*-deficient cells.

**Figure 3.**
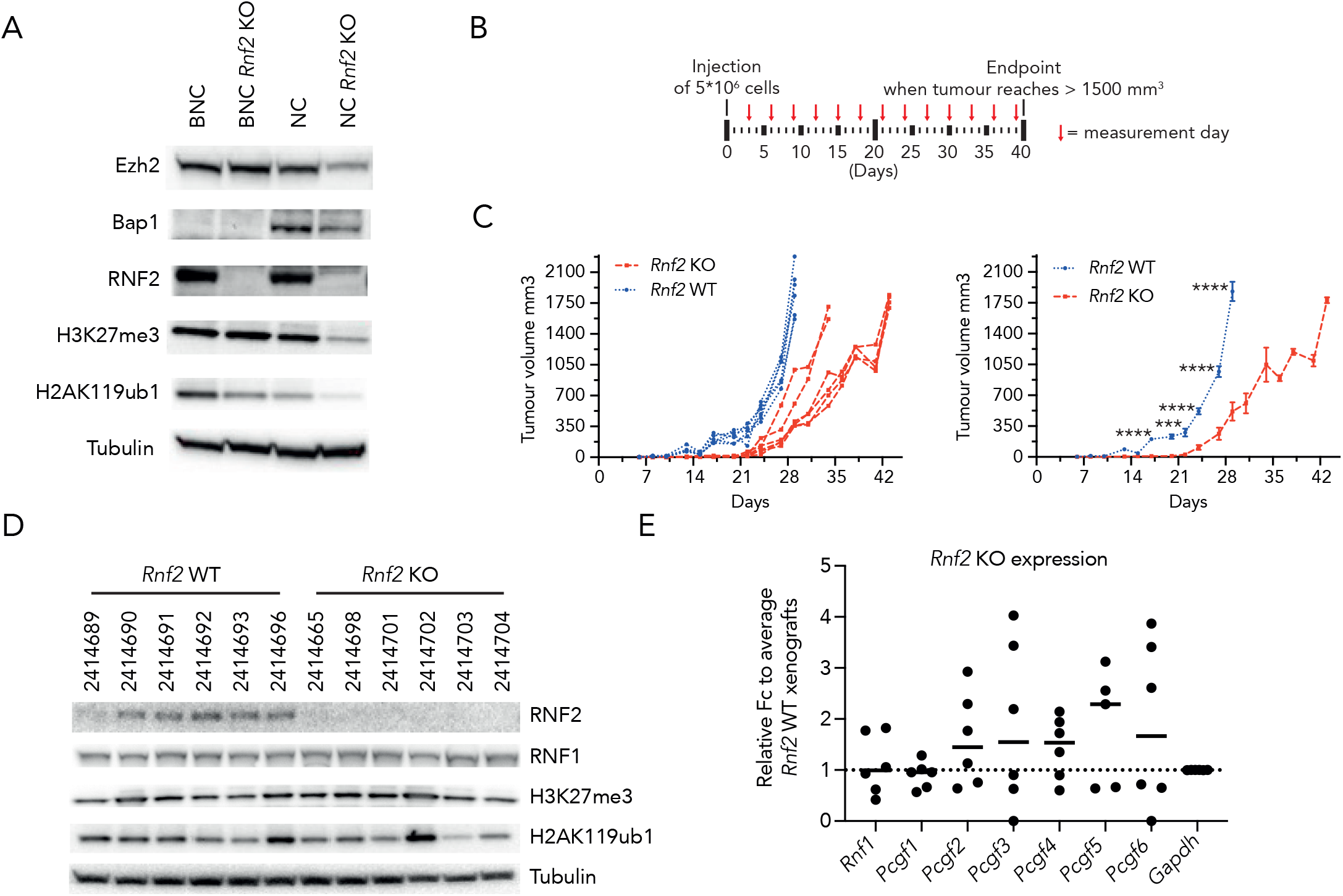
*Bap1*-deficient mesothelioma cells knocked-out for *Rnf2* have delayed tumour onset. **A**, Western blot of the BAP1 and RNF2 status of the cell lines, also showing the consequences of loss of these proteins on the PRC1 and PRC2 mediated chromatin marks. Tubulin was used as a loading control. **B**, Schematic overview of the xenograft experiments and the interval of tumour measurement. **C**, Tumour volume curves for the individual mice (left panel, n = 6) and the average values of the mice combined (right panel). P values were determined by two-tailed unpaired Student’s t-test; ***P < 0.001, and ****P < 0.0001. **D**, Western blot for the genetic status of the xenografts at the end of the experiment showing compensation on H2AK119ub1 in *Rnf2* knock-out tumours. **E**, qPCR data for fold changes of *Pcgf* genes in the *Rnf2* knockout tumours relative to *Rnf2* wild-type tumours showing an overall increase in expression (n = 6).

Next, we subcutaneously grafted *Rnf2* wild-type and *Rnf2* knock-out *Bap1*-deficient cells in the flanks of NOD-Scid IL2Rγnull mice (n = 6) and monitored the tumour outgrowth until the humane endpoint (**Figure 3b**). In concordance with the clonogenic potential of the cells *in vitro*, we observe that *Rnf2* knock-out cells grow out at a slower rate than *Rnf2* wild-type cells. On average *Rnf2* KO cells reached the maximum tumour volume 14 days later than the *Rnf2* WT counterpart, indicating that RNF2 is required for early tumour growth but that this can be overcome (**Figure 3c**).

To validate the genetic status of the tumours we performed western blot for RNF2 and H2AK119ub1. Unexpectedly and in contrast to the cells grown *in vitro*, H2AK119ub1 levels were not consistently lower in the *Rnf2* KO cohort suggesting that there might have been compensation for the loss of RNF2 (**Figure 3d**). As the PRC1 complex can rely on both RNF1 and RNF2 for its ubiquitination function we analysed whether RNF1 levels were compensating for RNF2, surprisingly we did not find consistently upregulated RNF1 neither on protein level nor on mRNA level (**Figure 3d and e**). In addition, we checked whether there might have been an upregulation of the PRC2-mediated H3K27me3 mark, however, we do not observe a difference in this mark between *Rnf2* KO and WT tumours (**Figure 3d**). To investigate whether there were changes in canonical and non-canonical PRC1 complexes we performed qPCR for the different *Pcgf* genes. Here we observe that in *Rnf2* knock-out cells, there is a trend towards upregulation of all the PCGF PRC1 subunits, with the exception of PCGF1 (**Figure 3e**).

Taken together, we show that loss of the E3 ubiquitin ligase RNF2 leads to a loss of clonogenic potential and a delay in tumour development in *Bap1*-deficient cells. In contrast, *Bap1*-proficient cells do not seem to be affected by loss of RNF2. *Rnf2* loss, *Bap1*-deficient cells eventually grow out to tumours and restore their H2AK119ub1 levels. As we see a trend in upregulation of PRC1 complex subunits in *Rnf2* knock-out cells it will be interesting to further investigate the role these subunits might play in the onset of *Bap1*-deficient mesothelioma.

## Discussion

The results in this chapter substantiate the important role that RNF2 and H2AK119 mono-ubiquitination have in the tumorigenesis and progression of *Bap1*-deficient mesothelioma. In accordance with a recent study in ES cells and our previously published data, we report an overall increase in the PRC1-mediated H2AK119ub1 mark upon loss of BAP1 (9, 10). We also described that loss of this mark, by knocking out the E3 ubiquitin ligase RNF2, leads to a reduced clonogenic capacity of *Bap1*-deficient mesothelioma cells but not that of *Bap1*-proficient cells. In addition, we show that this phenotype is also seen *in vivo* as a delay in tumour onset, however, these cells are still able to grow out to sizable tumours.

In mesothelioma it has been described that it is mainly the PRC2-mediated tri-methylation of H3K27 that determines the transcriptome in *Bap1*-deficient cells (10). BAP1 has been suggested to prevent chromatin compaction and the global redistribution of H3K27me3 (9). In addition, given the deubiquitinase function of BAP1 in the PR-DUB complex, it is not surprising that BAP1 also plays a role in regulating H2AK119 ubiquitination (12). Even though it is clear that all Polycomb members and their respective marks play a role in transcriptional regulation, their specific contributions remain unclear (15, 16, 17). Here we show, that despite the previously reported major role of H3K27me3, changes in H2AK119ub1 levels lead to a delay in tumour growth in *Bap1*-deficient mesothelioma. Together with the observations of limited dependency on PRC2 for PRC1 recruitment, this suggests that PRC1 is able to transcriptionally regulate a subset of genes independently of PRC2 (18, 19).

PRC1 complex exerts its ubiquitination function via the E3 ubiquitin ligases RNF1 and RNF2. Both RNF1 and RNF2 are important for cell viability and survival as they are key to the activity of canonical and non-canonical PRC1 sub-complexes (20, 21, 22, 23). Inactivation of *Rnf1* in mice leads to non-lethal developmental disorders (24), whereas *Rnf2* loss is embryonically lethal (25). However, there are many studies suggesting functional redundancy between RNF1 and RNF2 (26, 27). In this study we knocked-out *Rnf2* and showed that initial loss of H2AK119ub1 levels has been restored to wild-type levels in the injected tumour cell lines. This data might suggest compensation by the paralogue RNF1. However, even though RNF1 is still intact and able to form an active E3 ubiquitin complex, RNF1 mRNA and protein levels remained stable (26, 28). As H2AK119ub1 levels are restored over time but RNF1 is not upregulated we hypothesize that H2A ubiquitination in absence of RNF2 might be limited by the efficiency of the PRC1 complex with RNF1.

The PRC1 complexes are classified as canonical or non-canonical defined by the subunit interacting with RNF1/2, the PCGF proteins. These six different proteins (PCGF1 – 6) give rise to different PRC1 subcomplexes with varying properties (23, 29). Therefore, in this study we checked the relative abundance of the PCGF subunits between the *Rnf2* wild-type and knock-out tumours and observed that, with the exception of *Pcgf1*, all of these are upregulated in *Rnf2* knock-out tumours. This might suggest that loss of *Rnf2* is compensated via upregulation of these PCGF proteins to restore H2AK119ub1 levels and eventually tumour progression. It will be interesting to further delve into the role of the specific PRC1 sub-complexes in the development and progression of *Bap1*-mutant mesothelioma.

Overall, this report illustrates that, despite the major role of H3K27me3 in mesothelioma, the PRC1-mediated mark H2AK119ub1 is an important factor as well and deserving of more research.

### Limitations of the study

Here we show that loss of the E3 ubiquitin ligase RNF2 and subsequent loss of H2AK119ub1 leads to a delay in tumour progression indicating an important role for PRC1 in mesothelioma. However, the precise role of the different (non)-canonical variants of PRC1 remains an open question and is in need of further investigation. Also, here we studied the consequence of *Rnf2* loss in cell lines that were already malignant. To uncover the contribution of PRC1 to the development of mesothelioma, future work should focus on the consequence of *Rnf2* loss in autochthonous mouse models of mesothelioma, such as the models described by our lab previously.

## Materials and Methods

### Cell culture

Mouse mesothelioma cell lines with stable Cas9 were previously generated in our laboratory and cultured in Dulbecco’s Modified Eagle Medium/Nutrient Mixture F-12 (DMEM/F12+Glutamax; Gibco), supplemented with 4 μg/ml Hydrocortisone (Sigma), 5ng/ml murine EFG (Sigma), insulin-transferrin-selenium solution (ITS; Gibco), 10% fetal calf serum (FCS; Capricorn), and 1% penicillin and streptomycin (Gibco). Cell lines were maintained at 37 °C in a humidified atmosphere containing 5% carbon dioxide (CO2) and were tested for mycoplasma contamination using MycoAlert Mycoplasma detection kit (Lonza).

### Knock-out of RNF2 by CRISPR/Cas9

For RNF2 knock-out we used pLKO5.sgRNA.EFS.GFP (Addgene, 57822) vector targeting the following sequences: 5’-GAGTGGCGATTGATCCAGTCA-3’ and 5’-CCCTTAATTAGATTACAGCG-3’. Plasmids were transiently transfected into stably Cas9 expressing target cells using RNAiMax (Invitrogen, product #13778075). GFP positive cells were sorted by flow cytometry and sequenced by Sanger sequencing using the Mix2Seq kit (Eurofins).

### Cell growth assays

For proliferation rate assays cells were seeded in numbers that were determined prior to the experiment. Cells were counted with HyClone Trypan Blue (Cytiva) using a TC20 automated cell counter (Bio-Rad) and alive cells were seeded into 96-well plates in 100μl of culture medium and grown for 24, 72, and 120 hours. Cells were incubated for 4 hours with Resazurin (Sigma) and plates were read using an Infinite M1000 pro plate reader (TECAN). Cell proliferation was then calculated relative to control cell line.

For clonogenic assays, cells were seeded in 6-well culture plates and allowed to adhere overnight, 250 cells per plate. Cells were then refreshed with new medium every three days. After 10 days plates were fixed using 4% Paraformaldehyde (Merck) and stained with 0.1% crystal violet solution (Sigma) in PBS with 10% EtOH. Plates were analysed using the ImageJ plugin ‘Colony_Counter’.

### Western blot analysis

Whole-cell pellets were lysed in RIPA buffer (50 mM Tris, pH 8.0, 50 mM NaCl, 1.0% NP-40, 0.5% sodium deoxycholate, and 0.1% SDS) containing protease inhibitor cocktail (Complete; Roche) and phosphate inhibitors (10 mM NaF final concentration, 1 mM Na_3_VO_4_ final concentration, 25mM β-Glycerophosphate final concentration, 1mM PMSF, and 1 mM Na_4_P_2_O_7_ final concentration). Protein concentrations were measured using Protein Assay Dye reagent (Bio-rad) and a Nanodrop 2000c machine. Equal amounts of protein were loaded onto 4–12% Bis-Tris gels (NuPage-Novex, Invitrogen) and transferred onto nitrocellulose membranes (0.2 μm; Whatman). Membranes were blocked in 5% BSA in phosphate-buffered saline (PBS) with 0.1% Tween-20 (PBST) for 1 h, incubated with primary antibodies in PBST 1% BSA overnight at 4°C, and incubated with secondary antibodies coupled to HRP for 45 min in PBST 1% BSA at room temperature. Antibody detection was accomplished using Amersham ECL detection reagent (GE healthcare). Membranes were imaged on a BioRad ChemiDoc XRS+. The following antibodies were used for western blot analyses: BAP1 D7W70 (Cell Signalling, 13271S), Tri-Methyl-Histone H3 (Lys27) C36B11 (Cell Signalling, 9733S), anti-Tubulin (Sigma, T9026), DinG/RNF2 (gift from R. Kosecki), H2AK119ub1 D27C4 (Cell Signalling, 8240S).

### RT-qPCR

Extraction of RNA was done using ReliaPrep kit (Promega). Reverse transcription was performed with the Tetro cDNA synthesis kit (Meridian) making use of Random Hexamers. Power SYBR green master mix (Applied Biosystems) was used to perform qPCR in triplicates using the QuantStudio 5 Real-Time PCR System (ThermoFisher). Data were normalized against *Gapdh*.

### Animal studies

All animal procedures were performed in accordance with Dutch law and the institutional committees (Animal experimental committee and Animal welfare body) overseeing animal experiments at The Netherlands Cancer Institute, Amsterdam. Mice were housed under standard feeding, light cycles, and temperature with *ad libitum* access to food and water. All mice were housed in disposable cages in the laboratory animal center (LAC) of the NKI, minimizing the risk of cross-infection, improving ergonomics and obviating the need for a robotics infrastructure for cage-washing. The mice were kept under specific pathogen free (SPF) conditions.

To establish xenografts, 5x10^6^ mouse mesothelioma derived cells in 100μl PBS with 50% Matrigel (Corning) were subcutaneously implanted into the flank of 6 – 10 weeks old immune-deficient NOD-Scid IL2Rγnull (NSG) mice (Jackson Laboratory). Tumour growth was monitored by slide calliper 3 times a week (volume=length x width^2^/2). Mice were monitored daily for weight loss, signs of discomfort, abnormal behaviour, and death. Tumours were allowed to grow to ∼ 1500 mm^3^ in size before termination.

## Author Contributions

Conceptualization, N.L., J.B., M.v.L.; Methodology, N.L., D.H., J.B.; Formal Analysis, N.L., D.H.; Investigation, N.L., D.H.; Resources, J.B., M.v.L.; Writing, N.L.; Supervision, J.B., M.v.L.; Funding Acquisition, M.v.L.

## Acknowledgements

We thank the members of the animal facility of the Netherlands Cancer Institute for their support.

## Declaration of interests

The authors declare no conflict of interest.

## Notes

### Competing Interest Statement

The authors have declared no competing interest.

